# Structural and cellular insights into the inhibition of the drug efflux activity of the Hedgehog receptor PTCH1

**DOI:** 10.64898/2026.03.23.713596

**Authors:** Oussama Houha, Maryem Wachich, Claire Debarnot, Sandra Kovachka, Stéphane Azoulay, Isabelle Mus-Veteau, Valérie Biou

## Abstract

PTCH1, the receptor for the Sonic Hedgehog morphogen, mediates cholesterol transport across the plasma membrane by harnessing the proton motive force. In cancer, PTCH1 is frequently overexpressed and promotes chemoresistance by transporting drugs such as doxorubicin (dxr) out of cells. Among the inhibitors identified, PAH stands out for its ability to significantly enhance the efficacy of several chemotherapeutic drugs on melanoma and breast cancer cells.

To investigate PTCH1’s structure in complex with its inhibitor PAH, we overexpressed a construct spanning residues 1-619 and 720-1305 in HEK293 cells. The protein localized to the membrane, and transfected cells exhibited reduced sensitivity to dxr compared to control cells. Additionally, we observed a pH-dependent efflux of dxr, which was reversed by PAH, confirming that the PTCH1 construct used in this study functions as an active drug-efflux pump.

In the structure of PTCH1 bound to PAH determined using cryo-electron microscopy, PAH occupies a hydrophobic cavity in an extracellular domain which is normally occupied by cholesterol in other PTCH1 structures, and engages in a key hydrogen bond via one of its hydroxyl groups, a feature previously established as essential for its inhibitory function.

These findings not only clarify the molecular basis of PAH’s action but also provide a structural roadmap for rational drug design, enabling the development of next-generation inhibitors with enhanced potency.

## Introduction

### History of PATCHED as a HEDGEHOG receptor

The HEDGEHOG signaling pathway controls tissue patterning during the development of insects and vertebrates. After birth, it is involved in regeneration of many organs (such as bone, intestine, eye and brain) by controlling progenitor cell homeostasis. The HEDGEHOG morphogen (Sonic, Indian or Desert HEDGEHOG in mammals) is secreted by emitting cells as a dually-lipidated protein with a palmitate at the N-terminus and a cholesterol at the C-terminus ^1,2^.

PATCHED is a membrane bound protein that was identified as the HEDGEHOG receptor ^3^. It is a 150 kDa protein with 12 transmembrane helical domains, three cytoplasmic regions and two large, glycosylated, extracellular domains with three disulfide bridges ^4–9^. The three cytoplasmic regions are mostly predicted as disordered, and harbour two ubiquitylation sites associated with a shortened life-time of the protein ^10,11^. In the absence of HEDGEHOG, PATCHED inhibits another transmembrane protein of the G-Protein Coupled Receptor family, SMOOTHENED, that triggers a signalling cascade ending in target gene activation and cell differentiation. No stable protein-protein complex between PATCHED and SMOOTHENED has been observed. Instead, a messenger molecule was suggested to catalytically suppress SMOOTHENED activity ^12^. Due to the presence of a cholesterol at the C-terminus of HEDGEHOG morphogen ligand, a sterol-sensing domain (SSD) in the PATCHED transmembrane region, and of the similarity between PATCHED and the Niemann-Pick type C1 disease cholesterol transporter (NPC1), this messenger molecule could be a sterol-like compound ^12^. Indeed, our studies suggested that mammalian PATCHED (PTCH1) is responsible for cholesterol efflux from mouse fibroblasts, and that Sonic HEDGEHOG N-terminal domain (ShhN) binding decreases this efflux ^13^. This was confirmed when cryo-electron microscopy (cryo-EM) structures of PTCH1 showed several sterol-like densities ^4–9^. At least three sterol-binding sites were identified: at the SSD site, at the interface between the transmembrane and the extracellular domains (ECDs) forming a possible efflux pathway between the transmembrane domain and the extracellular matrix ^4,5,14,15^. Moreover, fully lipidated ShhN inserts its C-terminal cholesterol in the ECDs, slightly above the free cholesterol site, and its N-terminal palmitate is located at the membrane-ECD interface^15^.

### PATCHED, cancer and chemotherapy resistance

The HEDGEHOG pathway is tightly regulated both in development and stem-cell maintenance. Several genes involved in the HEDGEHOG pathway are mutated in cancer, amongst which *PATCHED1* (*PTCH1*) and *Suppressor of Fused*, two negative regulators whose inactivation increases HEDGEHOG signalling activity and cancer cell proliferation. Carcinoma from patients with Gorlin syndrome have been found to harbour mutations in the *PTCH1* gene, rendering the protein inactive and thus over-activating the HEDGEHOG pathway ^16,17^. In many cancers such as breast, lung, colon, prostate, pancreas cancers or melanoma, the Hh signalling pathway is aberrantly activated due to autocrine or paracrine production of Shh ^18–20^. PTCH1, being a Hh target gene, is over-expressed in many cancers and often associated with a bad prognosis ^21,22^

Moreover, abnormal Hh signaling activation and PTCH1 upregulation were reported in cells exhibiting resistance to chemotherapy, such as cancer stem cells or tumor-initiating cells ^23^. Elevated expression of PTCH1 was described in the residual gastric cancer cells of patients following chemotherapy treatment with cisplatin and was associated with poor survival of those patients^22,24^. Gonnissen et al. reported that upregulation of PTCH1 may be a prognostic marker for relapse in high-risk prostate cancer patients ^26^.

### PTCH1 is a drug efflux pump

Drug resistance mechanisms involved in cancer stem-cell persistence include drug efflux, epigenetic modifications, DNA repair increase, apoptosis inhibition ^27^.

Drug efflux in cancer cells is well-documented for the ATP-binding cassette transporter family, particularly P-glycoprotein. This family is so widespread that selectively inhibiting it remains a major challenge. Micro-RNA inhibition of the associated gene is being investigated as an alternative to drug treatment ^27,28^ .

PTCH1 presents structural similarities with the Niemann-Pick Type C protein 1 (NPC1) that imports cholesterol from lysosomal vesicles into the cytoplasm ^29^, and more generally with the efflux pumps from the resistance nodulation division (RND) family that use the proton motive force to expel antibiotics from bacteria ^30^.

Although PTCH1 is part of the RND family and is overexpressed at the surface of cancer cells, it is little studied as a cancer resistance factor. However, we showed for the first time that PTCH1 is a multidrug transporter involved in the resistance of cancer cells to chemotherapy ^31^. We expressed human PTCH1 in the plasma membrane of the yeast *Saccharomyces cerevisiae*, and we observed that the expression of PTCH1 conferred to yeast the ability to grow in the presence of chemotherapeutic agents such as doxorubicin (dxr), methotrexate or temozolomide. Our study showed that yeasts expressing PTCH1 were able to expel more dxr than control yeasts and that PTCH1 efflux activity uses the proton gradient as an energy source like the bacterial efflux pumps from the RND family ^31^. In more recent studies, we reported that the inhibition of endogenous PTCH1 expression in adrenocortical carcinoma (ACC) and melanoma cells using silencing RNA strongly decreased dxr efflux from these cells, indicating that PTCH1 contributes significantly to their chemotherapy resistance ^21,32^. Further evidence of the role of PTCH1 in chemotherapy resistance was provided by the observation that ACC and melanoma cells rendered resistant to dxr express more PTCH1 than parental cells. PTCH1 expels chemotherapy drugs from cancer cells by using the proton gradient made possible by the acidic extracellular environment – a hallmark of malignancy known as the Warburg effect, driven by cancer high glucose metabolism ^33^. Moreover, the HEDGEHOG signal is very low in adults, except in cases of stem cell maintenance, wound repair or neuro-regeneration. This makes PTCH1 a very attractive cancer specific drug-efflux target in adults and its inhibition should have few side effects.

### Attempts at reducing HEDGEHOG-related drug resistance

The screening of more than 2000 small molecules on the resistance of yeasts to dxr conferred by overexpressing human PTCH1 led us to the identification of a candidate inhibitor produced by a marine sponge: panicein-A hydroquinone (PAH). This molecule strongly inhibited the growth of yeasts overexpressing human PTCH1 in the presence of dxr ^34^. This effect was confirmed on melanoma cell lines endogenously expressing PTCH1, showing that PAH associated with dxr significantly decreased cell viability with respect to dxr alone ^34^. Furthermore, synthetic PAH showed an inhibition activity similar to that of the natural PAH, both against dxr and another drug, vemurafenib, in melanoma cells. Moreover, we showed that PAH enhances vemurafenib treatment efficacy on melanoma xenografts in mice without toxic effect ^32^. In a recent study, we reported that PAH also increased dxr and docetaxel efficacy on breast cancer cell lines ^35^.

In order to better understand the efflux inhibition activity of PAH towards PTCH1, we over-expressed PTCH1 in HEK293 cells, characterised its efflux activity and inhibition in cells, purified the protein and measured its affinity for PAH, and solved its cryo-EM structure in complex with PAH. We identified a PAH binding site and the residues involved in this binding pocket. These results provide key information on the mechanism by which PAH inhibits PTCH1 efflux activity.

## Results

### PTCH1 over-expressed in HEK293T cells has a drug efflux activity

Human Embryonic Kidney (HEK) 293T cells derived from HEK293 along with T-antigen of SV40 were transiently transfected with the pCAGEN vector containing the human PTCH1 cDNA (Uniprot entry Q13635) covering residues 1-619 and 720-1305 with an amino-terminal FLAG tag and a carboxy-terminal 10-His tag (pCAGEN-FLAG-PTCH1-10His). Western-blots with anti-FLAG antibodies confirmed the expression of the PTCH1 protein at the expected molecular weight (Fig. 1A). To obtain a stable cell line over-expressing PTCH1, HEK293T cells were co-transfected with the pCAGEN-FLAG-PTCH1-10His plasmid and a vector containing the blasticidin S deaminase (BSD) cDNA that confers resistance to blasticidin. Cells that integrated both plasmids were selected with medium containing increasing concentration of blasticidin, and finally stably cultured in medium containing 10 µg/mL blasticidin. The expression of PTCH1 in this stable cell line is close to that obtained after transient transfection (Fig. 1A). Immunofluorescence with anti-FLAG antibodies showed that PTCH1 is expressed at the plasma membrane in these cells (Fig. 1B). Remarkably, cells over-expressing PTCH1 grow differently than non-transfected HEK293T cells on plates. They form clones and spheroids (Fig. 1C), as previously observed in a small cell population endogenously over-expressing PTCH1 isolated from an adrenocortical carcinoma cell line ^24^. To facilitate the production of HEK293T cells stably expressing FLAG-PTCH1-10His (called HEK293T-PTCH1 below), these cells were adapted to grow in suspension using FreeStyle™ 293 medium containing 10 µg/mL Blasticidin.

**Figure 1.**
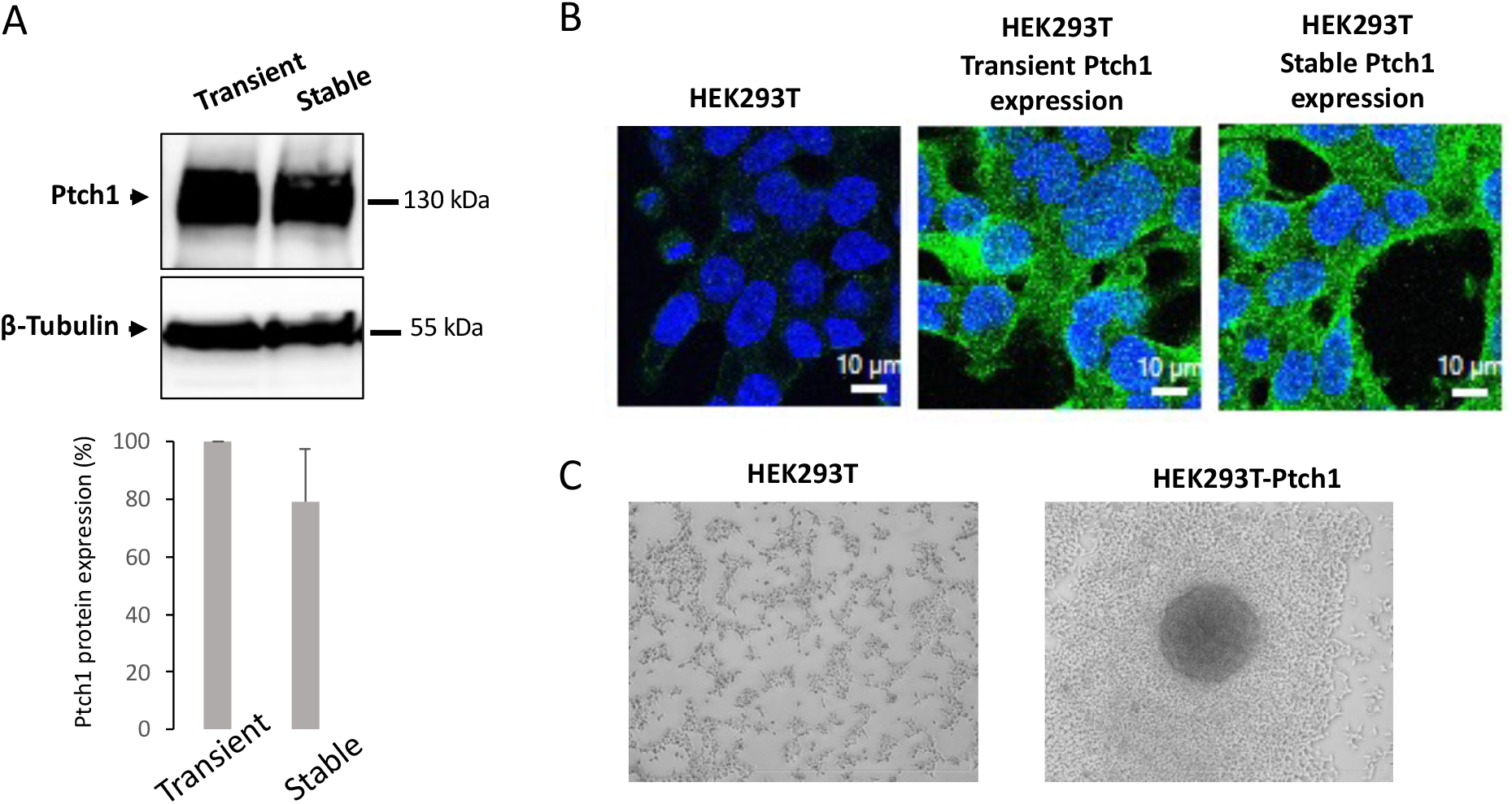
FLAG-PTCH1-10His is well expressed at the plasma membrane in HEK293T cells. (A) FLAG-PTCH1-10His expressed at similar levels in transiently transfected cells and in the selected stable cell line. Western blot analysis of cell extracts using anti-FLAG and anti-⍰-tubulin antibodies, and quantification of FLAG-PTCH1-10His expression relative to ⍰-tubulin using ImageJ software. (B) FLAG-PTCH1-10His localises to the plasma membrane in HEK293T cells for both transient and stable expression. Immunofluorescence labelling using anti-FLAG antibodies and FITC-coupled secondary anti-mouse antibodies (green). Images were acquired using a Leica SP5 confocal microscope with a 63× objective and 3× digital zoom. DAPI was used to stain the nuclei (blue). (C) Overexpression of Ptch1 modifies the growth of HEK293T cells. Images show HEK293T and HEK293T-PTCH1 cells seeded on plates.

We then evaluated the drug efflux activity of the PTCH1 construct to be purified and used for cryo-EM experiments using the stable cell line HEK293T-PTCH1.

HEK293T and HEK293T-PTCH1 cells at 80% confluency in 96 well plates were treated with increasing concentrations of doxorubicin (dxr) or docetaxel for 24 hours. Cell viability assays revealed that HEK293T-PTCH1 cells are less sensitive to these chemotherapeutic agents than control HEK293T cells, as demonstrated by a more than two-fold increase in the IC_30_ (the concentration required to kill 30% of cells) calculated from these assays (Fig. 2A and Table 1). From these experiments and the mean IC30 values, we then calculated the drug sensitivity score (DSS3) which is a systematic algorithmic solution for quantitative drug sensitivity scoring based on continuous modeling and integration of multiple dose-response relationships in high-throughput compound testing studies, and widely applicable to various experimental settings both in cancer cell line models and primary patient-derived cells ^36^ . Accordingly, PTCH1 expression reduces DSS3 by a factor of 5.9 for dxr and 1.75 for docetaxel (Fig. 2A). These results show that overexpression of FLAG-PTCH1-10His in HEK293T cells confers resistance to dxr and docetaxel.

**Figure 2.**
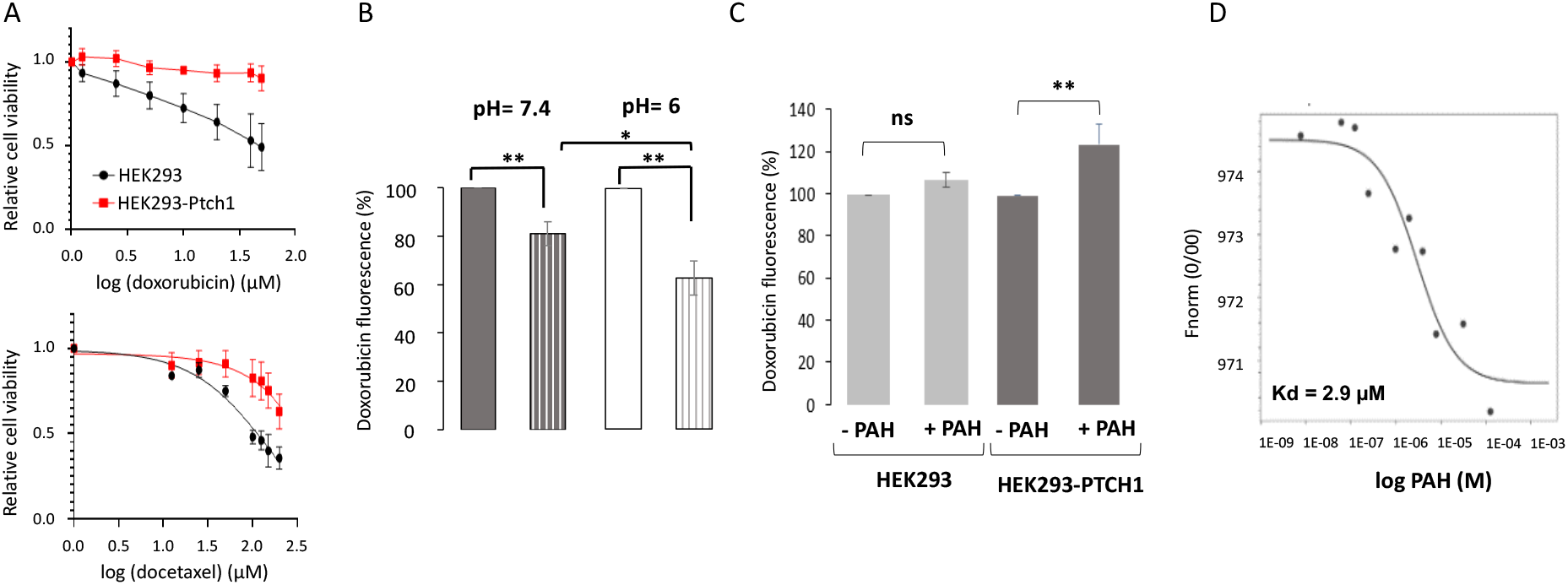
PTCH1 over-expressed in HEK293T cells exhibits drug efflux activity, which is inhibited by PAH. (A) FLAG-PTCH1-10His overexpression confers resistance to doxorubicin (dxr) and docetaxel to HEK293T cells. HEK293T cells (black) and HEK293T cells (red) stably expressing FLAG-PTCH1-10His were treated with increasing concentrations of dxr or docetaxel for 24 hours, after which cell viability was quantified using neutral red. (B) FLAG-PTCH1-10His stably overexpressed in HEK293T cells effluxes dxr using the proton gradient. The cells were incubated for 60 minutes with dxr at pH 7.4 or 6, after which they were fixed and the intracellular dxr fluorescence was quantified. Solid rectangles correspond to HEK293 and striped rectangles correspond to HEK293-PTCH1 cells. (C) PAH inhibits dxr efflux from HEK293T cells stably expressing FLAG-PTCH1-10His. Cells were incubated for 60 minutes with dxr at pH 6, either with or without PAH, and then fixed before the intracellular dxr fluorescence was quantified. **(D)** PAH binding to HEK293-PTCH1 membrane preparation. 30 µg/mL of membranes from HEK293-PTCH1 cells were incubated with tris-NTA-NT647 fluorescent probe, and then with 250 µM to 15 nM of PAH. After a short incubation, the samples were loaded into capillaries in a microscale thermophoresis (MST) analysis system (Monolith NT.115, LED:100% & MST: Medium). In the experiment reported, the calculated Kd was 2.9 µM. The average Kd calculated from four independent experiments is 2.8±0.8 µM.

**Table 1.** Quantification of doxorubicin and docetaxel sensitivity. IC30 values determined from at least 3 independent experiments using GraphPad and drug sensitivity score (DSS3.

|  | Doxorubicin |  | Docetaxel |  |
| --- | --- | --- | --- | --- |
| | IC30 (μM) | DSS3 (AU) | IC30 (μM) | DSS3 (AU) |
| HEK293 | $35 \pm 7$ | 18.2 | $56 \pm 9$ | 5.9 |
| HEK293-Ptch1 | >100 | 3.1 | $130 \pm 57$ | 1.75 |

The accumulation of dxr in HEK293T and HEK293T-PTCH1 cells after a one-hour incubation was then quantified. Results show that the dxr amount in HEK293T-PTCH1 cells is 81% of that in HEK293T cells, indicating that FLAG-PTCH1-10His over-expressed in HEK293T cells effluxes dxr (Fig. 2B). By lowering the pH of the buffer to 6, the amount of dxr in HEK293T-PTCH1 cells is 63% that in HEK293T cells while the amount in HEK293T cells remains similar at both pH values, indicating that PTCH1 over-expressed in HEK293T cells effluxes more dxr at lower external pH and uses the proton motive force to efflux dxr as previously shown ^31^. Moreover, adding the PTCH1 drug efflux inhibitor, PAH, during the incubation with dxr increases the amount of dxr in HEK293T-PTCH1 cells by 21.7±5.7 % (Fig. 2C), indicating that PAH interacts with PTCH1 over-expressed in HEK293T cells and inhibits its efflux activity as shown in ^32^. Binding experiments on membranes prepared from HEK293T-PTCH1 cells using microscale thermophoresis ^37^ allowed us to estimate that PAH interacts with PTCH1 over-expressed in HEK293T cells with an affinity of 2.8±0.8 µM (Fig. 2D). This affinity is in the same order of magnitude as that previously published on full-length PTCH1 expressed in yeast ^32^.

Overall, these experiments further confirm that the PTCH1 protein over-expressed in HEK293T cells is able to efflux chemotherapeutic drugs such as dxr or docetaxel, and that PAH inhibits its efflux activity, as shown on the wild-type PTCH1 expressed in yeasts and endogenous PTCH1 in melanoma cells ^32^. PTCH1 construct spanning residues 1-619 and 720-1305 over-expressed in HEK293T cells therefore exhibits the same properties as the wild-type endogenous PTCH1, and can be purified and used for biochemical and structural studies of its interaction with PAH.

### Structure of PTCH1 bound to the efflux inhibitor PAH

Because of the scalability of their ability to grow in suspension in a bioreactor, HEK-ExpiF293 cells transiently transfected with the pCAGEN-FLAG-PTCH1-10His plasmid were used to produce FLAG-PTCH1-10His (1-619 and 720-1305) for purification, affinity measurement and cryo-EM structural studies.

PTCH1 was solubilised using dodecyl ß-maltoside (DDM) and cholesterol hemisuccinate (CHS), and purified using anti-Flag affinity resin in the absence of CHS. Figure 3A shows the SDS-PAGE of the elution fractions with a major band above the 140kDa marker, that was identified as PTCH1 by mass spectrometry (not shown).

**Figure 3.**
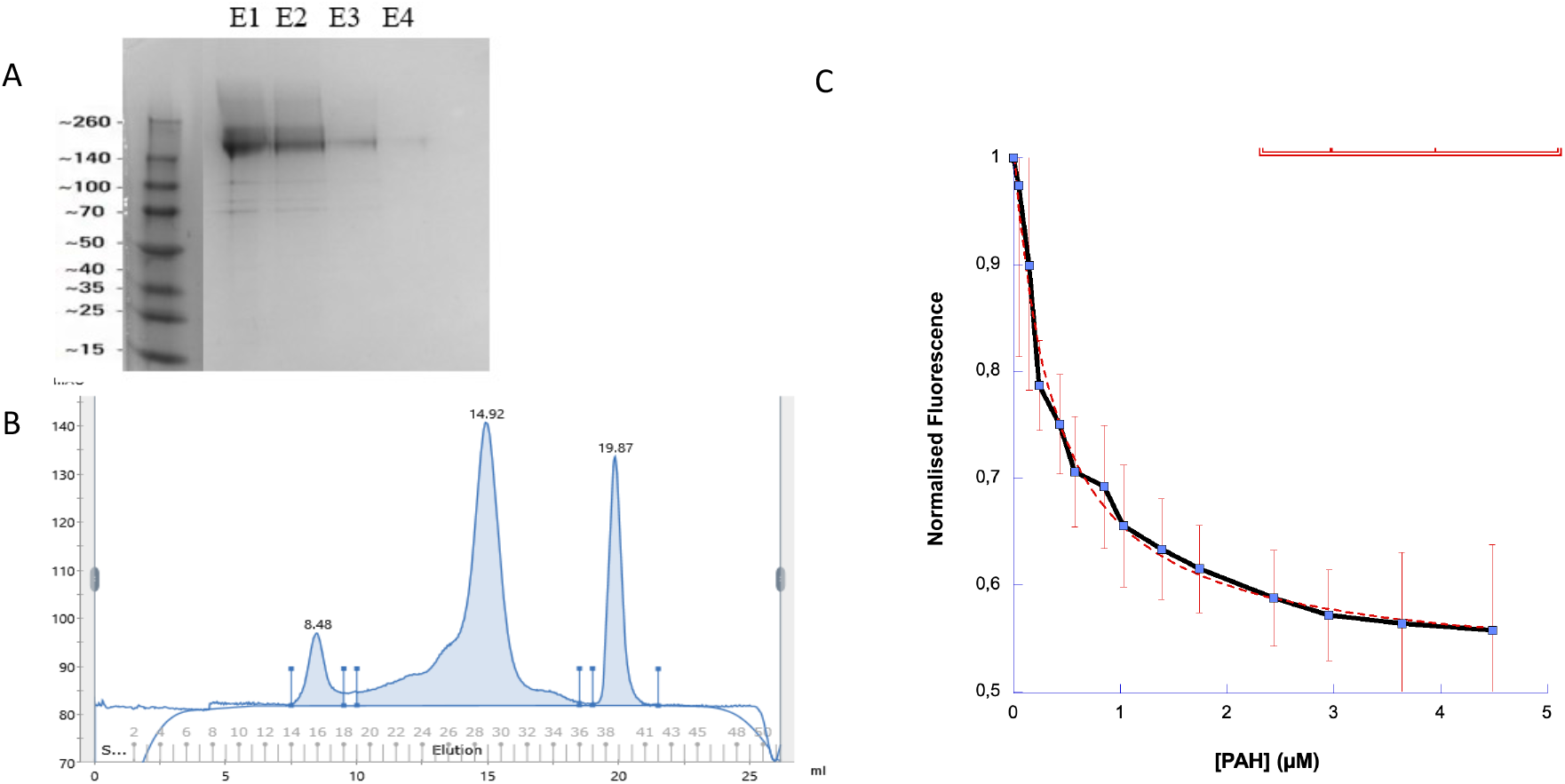
PTCH1 purification and affinity measurement for PAH. (A) Coomassie Blue stained SDS-PAGE 12% acrylamide gel: elution fractions E1, E2, E3, and E4. (**B**) Superose 6 10/300 SEC chromatogram: the central peak at 14.9 mL corresponds to the PTCH1 protein, the first peak corresponds to aggregates, and the last peak corresponds to the Flag peptide. (C) Affinity measurement using normalised tryptophan fluorescence as a function of PAH concentration.

We then measured the affinity of PAH for purified PTCH1 using tryptophan fluorescence. Flag-PTCH1-10His construct contains 19 Trp residues. We observed that the addition of increasing concentrations of PAH to purified PTCH1 triggered a decrease in the intrinsic fluorescence signal, and the experimental curve can be fitted with a hyperbola indicating a single binding site with a dissociation constant of 0.376 ± 0.046 µM (figure 3C). This shows that PAH directly binds PTCH1 purified in detergent.

To solve the cryo-EM structure of the PTCH1:PAH complex, the elution fractions were concentrated and injected in a size-exclusion column in a buffer containing 0.0075% LMNG. The chromatogram starts with a minor aggregation fraction, followed by a main peak at 14.9 mL that contains PTCH1 and a second peak at 19.9 mL corresponding to the Flag peptide (Figure 3B). The main peak fractions were pooled and concentrated to 1 mg/mL and incubated with PAH at a protein:PAH molar ratio of 1:50 for 30 minutes on ice before being deposited on cryo-EM UltraAuFoil grids and vitrified. As described above, we collected 25810 movies which were processed according to the scheme in supplementary figure S1. The best map resulted from 79080 particles with a 3.4Å resolution (Figure 4A and Table 2). The cryo-EM model of PTCH1 bound to PAH inhibitor protein model spans residues 73 to 607 and 731 to 1185 and is very similar to that of other human PTCH1 structures bound to cholesterol hemisuccinate. It shows a 12-helix transmembrane region and two amphipathic helices at the cytoplasm-membrane interface, with a pseudo two-fold symmetry. Two extracellular domains ECD1 and ECD2 present four glycosylation sites and three disulfide bridges (fig. 4A).

**Table 2.**
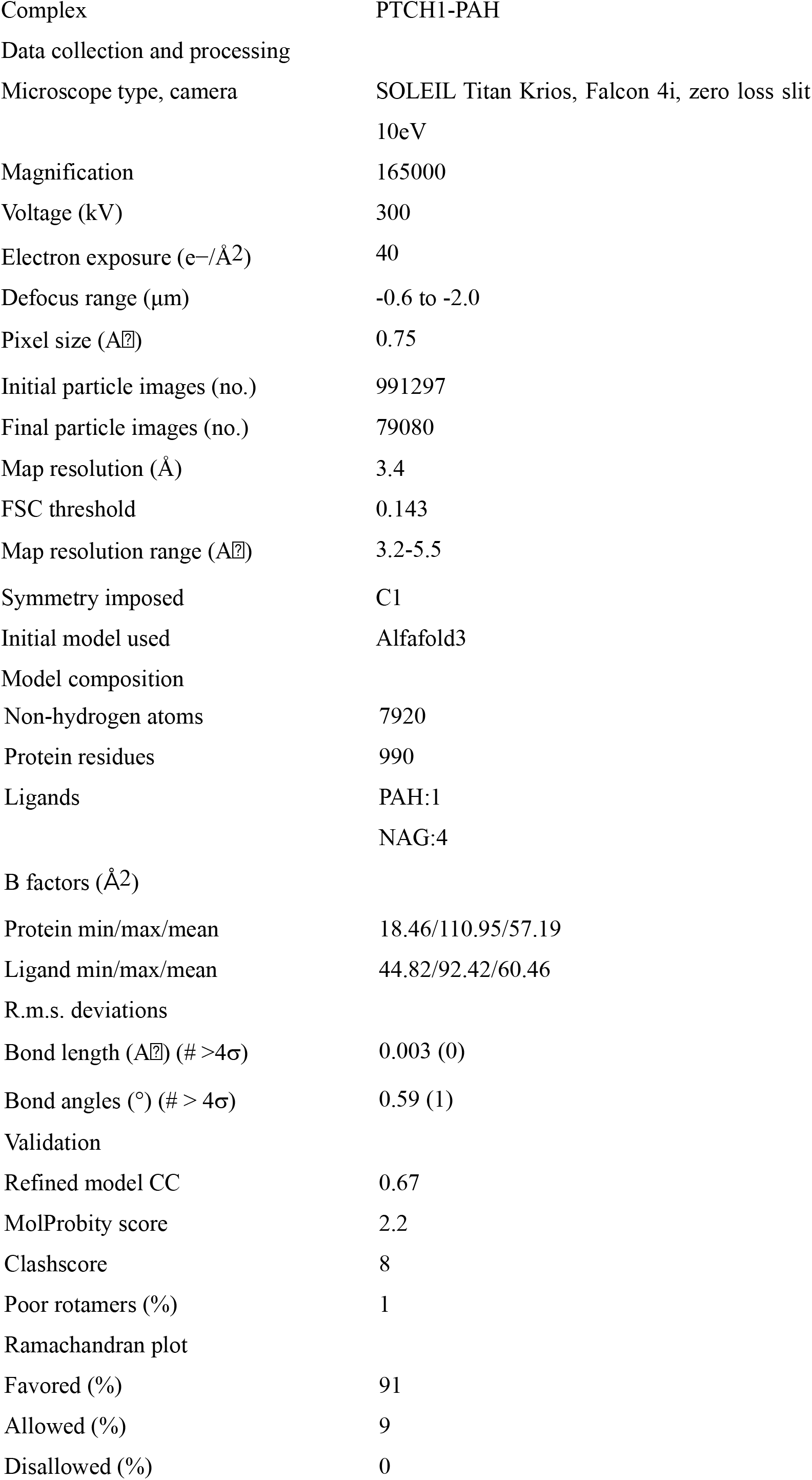
Cryo-EM Data collection and model refinement

**Figure 4.**
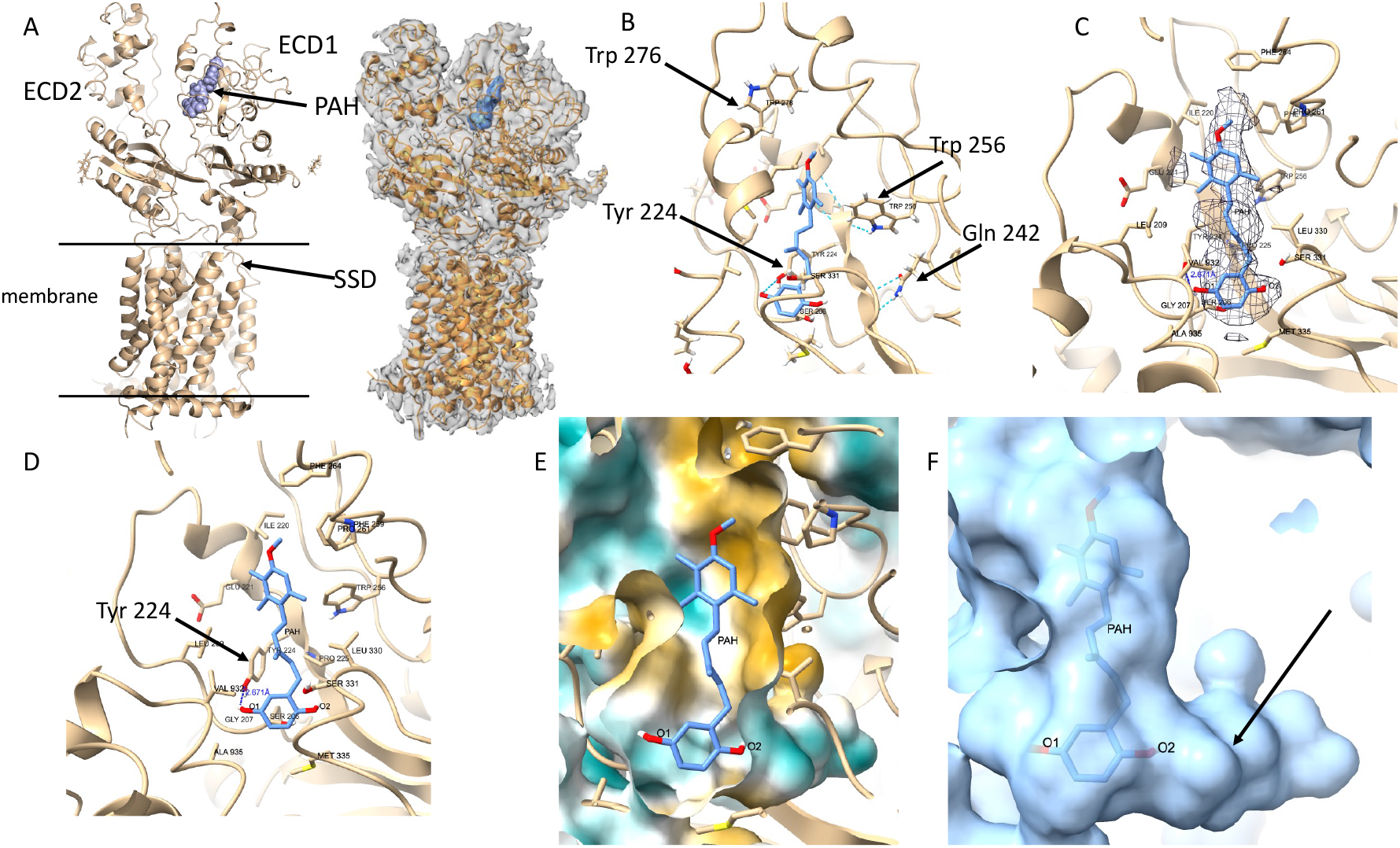
Cryo-EM structure of the PTCH1:PAH complex. A, overall structure of PTCH1 with the extracellular domains ECD1 and ECD2, the PAH location, the sterol sensing domain SSD and the trans membrane area. B, detailed view of the PAH binding site with residues mentioned in the text. the dashed lines indicate hydrogen bonds. C, PAH binding site with Coulomb density in the PAH zone. D, same view without the density, showing the H-bond between PAH O1 and Tyr 224 hydroxyl functions. E, same view with the protein surface colored as a function of hydrophobicity. Orange is hydrophobic and blue is hydrophilic. F, surface representation of the protein surface showing the druggable pocket near PAH O2 hydroxyl marked by a black arrow.

The transmembrane region shows a fold similar to the other PTCH1 structures, for example 6DMB ^5^. We don’t notice any ligand density in the sterol sensing domain between helices TM1 and TM2, indicating that all of the CHS has been eliminated during purification. Instead, residues 115-118 at the C-terminus of TM1 helix diverge from helical structure to form a bend towards the CHS binding site observed in other structures (Fig. 4A and Fig. 5C-D).

**Figure 5.**
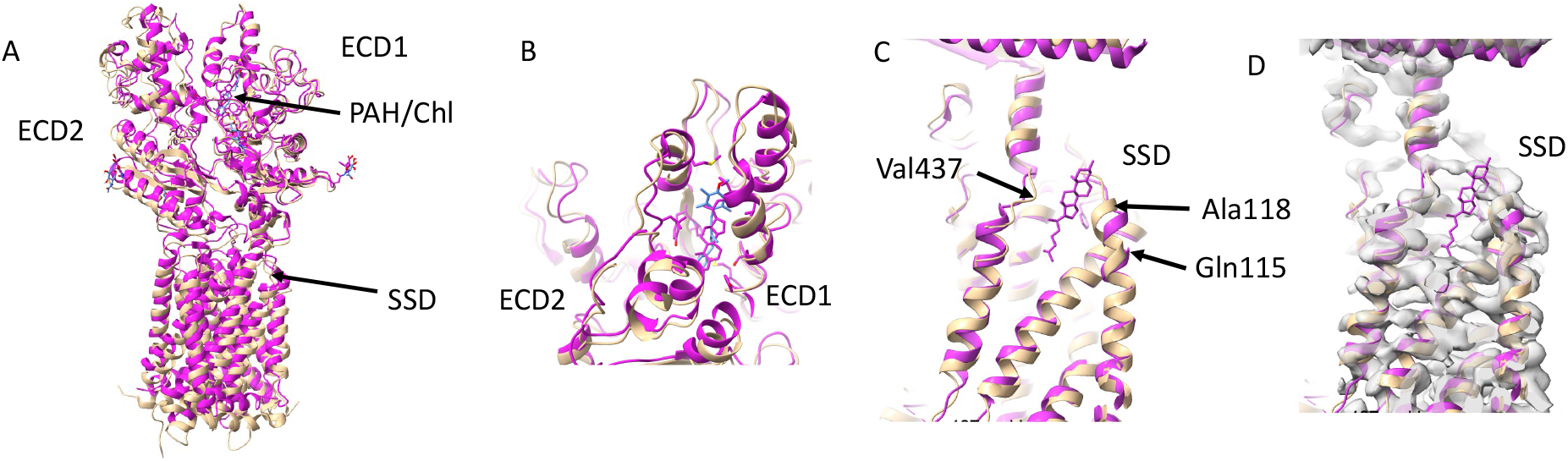
PAH binds in the same site as cholesterol in the ECD region and the sterol sensing domain is empty. All figures feature superimposition of 6RVD chain B (magenta) and this structure (beige). (A) global view indicating the PAH binding site and SSD motif. (B) Focus on the PAH binding site shows that PAH and cholesterol bind the same site. (C) In the SSD region shows a distortion of TM1 and TM2 helices closing the sterol binding site. D, same figure with cryo-EM map from Ptch1-PAH complex.

A non-protein cryo-EM density is visible in the ECD1 close to its interface with ECD2, in an aromatic-rich site containing Tyr 224, Trp 256, Phe 259, Phe 264, Trp 278 (Fig. 4B). It easily accommodates a PAH molecule. The proximity of the two tryptophan residues to the PAH site explains the rearrangement that induces the fluorescence change we observed upon PAH binding (Fig. 4B). The PAH is surrounded by two short helices (residues 213-222 and 277-285) and loops without a regular secondary structure (Fig. 4B). The density map shows that the PAH exhibits an elongated conformation (fig 4C-D) with the trimethyl-anisole ring close to several aromatic and hydrophobic residues and its methoxy moiety is accessible to the solvent. The hydroquinone moiety is located in a buried pocket and its hydroxyl in *meta* forms a hydrogen bond with the Tyr 224 side chain hydroxyl, thus stabilising the interaction with the protein (see O1 oxygen in Fig 4B-C). Tyr 224 is located on a beta-strand region stabilised by hydrogen bonds between residues 222-226 and neighbouring residues Gln 242 and Trp 256 (Fig 4D). Fig 4E shows the protein surface coloured as a function of hydrophilicity according to the Kyte-Doolittle scale ^38^. The trimethyl-anisole ring region binding is mostly hydrophobic (orange) and the hydroquinone region is more polar (blue), making the interaction with the hydroquinone more stable (Fig. 4E). The *ortho* hydroxyl O2 is in a hydrophilic region that forms a large pocket as shown by fig 4F.

## Discussion

### PTCH1 overexpressed in HEK293 cells is functional in mediating the efflux of chemotherapeutic drugs

We generated a stable cell line expressing levels of PTCH1 equivalent to those in transiently transfected cells. In both cases, PTCH1 was visualized at the cell membrane, confirming its proper folding. Drug efflux was monitored and found to be more efficient at pH 6 than at pH 7.4, indicating that PTCH1 uses the proton motive force for its transport activity. Additionally, treatment with the PAH molecule very significantly increased intracellular dxr levels in HEK293-PTCH1 cells compared to non-transfected HEK293, confirming its inhibitory effect on PTCH1.

A drug efflux channel is blocked via polar and hydrophobic interactions between PTCH1 and PAH. We solved the cryo-EM structure of the PTCH1-PAH complex. The protein conformation is very close to that of the cholesterol-bound state, for example in the 6RVD structure ^15^ (the rmsd between both ECD1 structures is 1.8Å over 161 residues) (Fig 5A and B). In this structure, the PAH binding site to PTCH1 is located in the ECD1 at the interface with ECD2, occupying a region where cholesterol is found in other structures of PTCH1. Within this site, the trimethyl anisole ring of PAH is embedded in an aromatic rich region and the hydroquinone engages in more polar interactions with the protein, compared to cholesterol or cholesterol hemisuccinate, cholesterol being a transport substrate and cholesterol hemisuccinate an analogue of cholesterol.

The binding site lies along a channel going from the SSD to the extracellular region of PTCH1 (fig 6A) and the PAH effectively blocks this channel. Two additional cholesterol (or CHS) binding sites were identified in PTCH1 cryo-EM structures published previously: at the SSD and at the junction between the membrane and ECD region.

**Figure 6.**
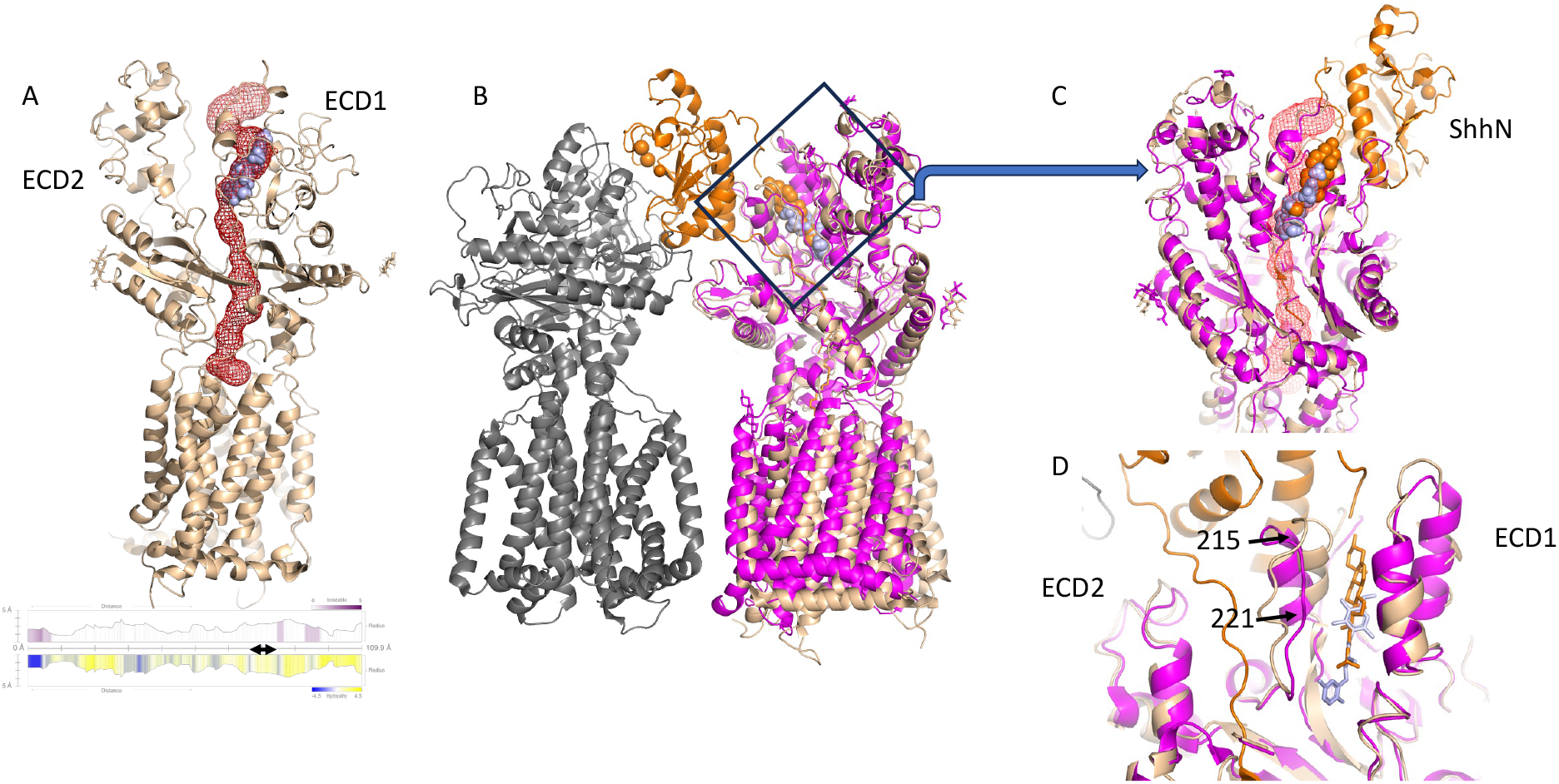
The channel of Ptch1 is blocked by PAH and its ligand ShhN. (A, Top) The channel detected by the Mole software (red mesh) is blocked by PAH (light blue). Below, the channel profile is displayed as a function of ionizability (top) and hydropathy (bottom). The double arrow indicates the PAH binding site. (B) Superimposition of the structure of Ptch1/PAH complex (beige with light blue PAH) and the structure of Ptch1/ShhN complex (6RVD with Ptch1 chain A magenta and chain B grey, SONIC HEDGEHOG and cholesterol orange). The frame shows the approximate location of fig 6CB. (D) same superimposition with a close-up view of the PAH binding site and the channel. Helix 215 to 221 is more open in 6RVD due to the cholesterol presence. C, same superimposition of the structure of Ptch1/PAH complex shows that the channel is blocked by both ligands.

This experimentally defined ECD1 binding mode indicates that PAH can occupy a bona fide sterol-binding pocket in the extracellular domain, rather than binding preferentially to the membrane-ECD “Neck” pocket as suggested by previous docking-based models ^39^.

This single binding site agrees with the fact that the affinity measurement curves measured on both the membranes and the purified PTCH1 protein are compatible with a PTCH1:PAH stoichiometry of 1:1 (Fig 2D and 3C).

Comparison with the structure of the complex between fully lipidated Sonic Hedgehog (ShhN) and PTCH1 (PDB id 6N7H)^8^ reveals that, while the overall structure of PTCH1 is similar, the membrane region shifts when the ECD domains are used as a reference for superimposition (Fig 6B). In addition, the cholesterol (CLR) covalently bound to the C-terminal of ShhN is located in a site adjacent to that of PAH, closer to the protein surface (Fig. 6C). As a result, the centres of mass of CLR and PAH are about 8.5Å apart within the same channel (Fig 6B-C). The channel provides an anchor point for ShhN palmitate or cholesterol moiety when it interacts with PTCH1, leading to a potential inhibition of cholesterol efflux and subsequent internalisation of the complex.

The cryo-EM structure shows that the hydroquinone moiety of PAH is positioned in a well-defined, buried subpocket that is amenable to structure-based ligand optimization. The meta-hydroxyl group of the hydroquinone forms a direct hydrogen bond with the side-chain hydroxyl of Tyr224, providing a strong directional anchor that rationalizes previous structure–activity relationship data showing that replacement of this meta-hydroxyl by an O-methyl abolishes inhibitory activity^32^. In line with this, computational analyses of PAH analogues on PTCH1 indicate that active compounds share a hydroquinone-like aromatic ring bearing hydrogen-bond donor groups and establish frequent contacts with residues such as Tyr224 within the ECD1 tunnel region ^39^, whereas inactive analogues lacking these features preferentially adopt self-associated, closed conformations and show less favorable interaction patterns. By contrast, the ortho-hydroxyl group projects toward a spacious, hydrophilic pocket, suggesting a natural “growth vector” for medicinal chemistry: increasing the size and polarity or introducing heteroatom-bearing substituents at the ortho position could exploit this cavity to enhance both binding affinity and target selectivity.

In addition, the trimethyl-anisole moiety constitutes a complementary handle for optimization, as its aromatic core is tightly packed against a predominantly lipophilic subpocket lined by hydrophobic and aromatic side chains, while the methoxy group projects toward the solvent and can be modified to modulate overall lipophilicity, target selectivity, or pharmacokinetic properties without perturbing the key hydrophobic contacts that appear to underlie high-affinity binding.

Comparison with structures of RND transporter proteins in complex with their inhibitors Several structures of RND transporters bound to transport inhibitors have been solved. For example, the structure of the bacterial antibiotic resistance protein AcrB in complex with the ABI-PP inhibitor shows that it binds to a hydrophobic pocket in the periplasmic region ^40^. In this case, the transport mechanism depends on allosteric changes between the three monomers of AcrB and the inhibitor blocks the conformation changes, thus preventing substrate exit. The structure of the cholesterol transporter NPC1 shows itraconazole inhibitor bound to a pocket at the junction between the membrane and the cytoplasmic region, near the binding site of the palmitate N-terminal modification of ShhN on PTCH1 ^41^ . NCR1 is an NPC1 homolog from *Plasmodium falciparum* that transports cholesterol from the host cell, and is also inhibited by compounds binding at the interface between the membrane region and the extracellular domains ^42^.

Therefore, whereas inhibition of bacterial RND transporters is predominantly achieved through allosteric modulation, inhibition of human RND drug efflux transporters such as PTCH1 and NPC1 appears to rely on ligand binding within the efflux pathway itself.

In summary, this study provides a comprehensive structural and functional characterization of the mechanism of PTCH1-mediated drug efflux inhibition by PAH. The data confirm our previous study ^31^ showing that PTCH1 utilizes the proton motive force for transport, and reveal that PAH effectively binds in the efflux channel through polar and hydrophobic interactions. This structure demonstrates that PTCH1 drug efflux inhibition occurs directly in the efflux channel as previously reported for other cholesterol transporters like NPC1. These insights not only deepen our understanding of PTCH1 inhibition mechanics but also pave the way for the rational design of novel inhibitors, with potential applications in overcoming drug resistance and improving therapeutic efficacy.

## Materials and methods

### Reagent preparation and storage

HEK293T and ExpiF293 cells were obtained from ATCC. Doxorubicin (MedChemExpress) was prepared in Milli-Q water at a stock concentration of 10 mM and stored at −20 °C, protected from light. Docetaxel (ThermoFisherSci) was prepared in DMSO at a stock concentration of 50 mM and stored at −20 °C.

Panicein A hydroquinone (PAH) synthesis was revised compared to the method published by Signetti et al. in 2020 ^32^ in order to selectively synthesise the E-PAH naturally occurring form (Kovachka et al, in preparation). PAH was prepared in DMSO at a stock concentration of 50 mM and stored at −20 °C. Construct, cloning, vector

The cDNA of human PTCH1 corresponding to Uniprot entry Q13635 covering residues 1-619 and 720-1305 was cloned into the pCAGEN (Addgene) vector with an amino-terminal FLAG tag and a carboxy-terminal His10 tag.

### Adherent cell culture

HEK293T cells were cultured in Dulbecco’s Modified Eagle Medium (DMEM) GlutaMAX™ (Gibco), supplemented with 10% (v/v) heat-inactivated fetal bovine serum. Cells were passaged using 0.25% trypsin-EDTA (Gibco).

### Generation of cell lines stably expressing FLAG-PTCH1-10His

HEK293T cells were co-transfected in 6-well plates with the pCAGEN-FLAG-PTCH1-10His plasmid and the pcDNA™6/V5-His A vector (Invitrogen™) containing the blasticidin S deaminase (BSD) cDNA that confers resistance to blasticidin at a 1:1 ratio using the jetPRIME® transfection reagent (Polyplus), according to the manufacturer’s instructions. Each transfection was performed using a total of 2 µg of DNA per well. 48 hours post-transfection, cells were selected with medium containing 3 µg/mL blasticidin for 72 hours. During this period, the culture medium was replaced once to remove dead cells and allow the surviving cells to reach approximately 80–90% confluence prior to passaging. Once split as previously described, half the cells were maintained in medium containing 3 µg/mL blasticidin, while the blasticidin concentration for the remaining cells was increased to 6 µg/mL. This selection procedure was repeated until cells were stably cultured in medium containing 10 µg/mL blasticidin.

### Suspension adaption

HEK293T cells stably expressing FLAG-PTCH1-10His (HEK293T-PTCH1) were seeded in culture flasks and initially maintained in 100% DMEM GlutaMAX™ supplemented with 10% (v/v) heat-inactivated fetal bovine serum and 10 µg/mL blasticidin, at 37°C in an 8% CO_2_ atmosphere under agitation at 150 rpm. Cells were passaged once a week. At each passage, cells were split, and FreeStyle™ 293 medium (Gibco) was gradually introduced, increasing from 25% to 100% of the total medium. The blasticidin concentration was maintained at 10 µg/mL throughout the adaptation process and subsequent cell culture.

### Western Blot

Cell lysis for all tested cell lines was performed on ice using RIPA buffer (50 mM Tris-HCl pH 8, 150 mM NaCl, 1% NP-40, 0.5% sodium deoxycholate, 0.1% SDS, 5 mM EDTA pH 8) supplemented with a protease inhibitor cocktail (cOmplete™, Roche) and 1 mM phenylmethylsulfonyl fluoride (PMSF). The concentration of proteins in the cell extracts was determined using Bradford protein assay (Bio-Rad). A total of 70 µg of protein per sample was separated by SDS-PAGE on an 8% acrylamide gel using Tris-SDS running buffer. Proteins were transferred onto a nitrocellulose membrane for Western blot analysis. After 1 h at room temperature in blocking buffer (PBS, 0.1 % Tween-20, and 5 % non-fat milk), membranes were incubated overnight with mouse anti-FLAG antibodies (ANTI-FLAG® M2 Sigma-Aldrich, diluted by 1/1000) and rabbit anti-GAPDH antibodies (Elabscience,1/20000). After 3 washes, membranes were incubated for 45 min with anti-rabbit (1/2000) or anti-mouse (1/5000) immunoglobulins coupled to horseradish peroxidase (Dako). Detection was carried out with an ECL Prime Western Blotting detection reagent (SuperSignal West Femto Maximum Sensitivity Substrate from ThermoScientific) on a Fusion FX imager (Vilber Lourmat).

#### Immunofluorescence

Cells were seeded on coverslips in a 24 wells plate. At 80% confluency, cells were washed twice with PBS, fixed with 4% paraformaldehyde (PFA) in phosphate-buffered saline (PBS) for 10-15 mins, and permeabilized with PBS/Triton 0.1% for 10 mins with agitation twice. For blocking, cells were incubated with 2% BSA in PBS for 30 mins with agitation. Coverslips were then incubated with anti-FLAG antibodies PBS/0.1% BSA (ANTI-FLAG® M2 Sigma-Aldrich, 1/1000) ON at 4°C. After 3 washes with PBS/0.1% BSA, coverslips were incubated with an anti-mouse antibody coupled to FITC (1/400) at RT for 45 mins. After 3 washes with PBS/BSA 0.1%, coverslips were mounted in glass slides with SlowFade Diamond Antifade mountant containing DAPI (Invitrogen). Images were acquired using a Leica SP5 confocal microscope with a 63X objective.

### Cytotoxicity assays

HEK293T and HEK293T-PTCH1 cells were seeded in 96-well plates in DMEM GlutaMAX™ supplemented with 10% (v/v) heat-inactivated fetal bovine serum (and 10 µg/mL Blasticidin for HEK293T-PTCH1 cells), and allowed to grow until reaching approximately 80% confluence.

Cells were treated with 0, 1.25, 2.5, 5, 10, 20, 40, and 50 µM of dxr or 0, 12.5, 25, 50, 100, 125, 150, 200 µM of docetaxel for 24 hours at 37°C in a 5% CO_2_ atmosphere. Dxr and docetaxel dilutions were prepared in DMEM GlutaMAX™ supplemented with 10% (v/v) heat-inactivated fetal bovine serum as described previously^32^. At the end of the incubation period, the treatment medium was removed, and cells were incubated for 3 hours at 37°C with 100 µL/well of neutral red (NR) solution (50 µg/mL in medium) to allow NR uptake by lysosomes in living cells. The NR solution was then removed, and the retained dye was released using a revelation solution (50% ethanol, 49% Milli-Q water, and 1% acetic acid). Absorbance measurements were performed using microplate readers (Multiskan Go Microplate Spectrophotometer, Thermo Scientific, US; and SPECTRA, Tecan, Männedorf, Switzerland). The concentration corresponding to y= 0.70, also known as IC30 (the concentration resulting on 30% cell death) is obtained by linear interpolation using the relation: IC30= (0.70 - *b*)/*a* where *a* is the slope of the straight line between two points A (x_1_, y_1_) and B (x_2_, y_2_) so that y_1_ < 0.70 < y_2_, and *b* is the intercept. Drug Sensitivity Scores (DSS3) were calculated in R (version 4.5.1) using the *drc* and *MESS* packages, as previously described by ^36^ .

### Doxorubicin accumulation measurements by flow cytometry

HEK293T and HEK293T-PTCH1 were detached using Accutase® solution (Sigma) prior to cell counting. For each pH condition, 5 × 10^5^ cells were used. Cells were then centrifuged, and the pellet was washed once with physiological buffer (140 mM NaCl, 5 mM KCl, 1 mM CaCl_2_, 1 mM MgSO_4_, 5 mM D-glucose, 20 mM HEPES) adjusted to either pH 6.0 or pH 7.4. Afterwards, cells were incubated for 1 hour with 5 μM dxr prepared in the corresponding pH-adjusted buffer. Untreated HEK293T cells were used as negative controls to assess cellular auto fluorescence. After incubation, dxr solutions were discarded, and cells were fixed for 10 minutes at 4 °C using 4%PFA in phosphate-buffered saline (PBS). Cells were then washed twice with PBS prior to signal acquisition. DAPI (3 μL at 5 mg/mL) was added to each sample to discriminate dead cells. Flow cytometry analysis was performed within 1 hour using a BD LSR II Fortessa flow cytometer (BD Biosciences).

### Doxorubicin efflux inhibition measurements by fluorescent microscopy

HEK293T and HEK293T-PTCH1 cells were seeded on coverslips in 24-well plates and allowed to reach approximately 80% confluence. Cells were washed once with physiological buffer (140 mM NaCl, 5 mM KCl, 1 mM CaCl_2_, 1 mM MgSO_4_, 5 mM D-glucose, 20 mM HEPES, pH 6.0) before treatment either with 5 μM dxr or with 50 µM PAH for 5 minutes before adding 5 μM dxr. Cells were then incubated for 1 hour at 37°C and 5% CO_2_, and protected from light. Afterwards, coverslips were fixed with 4% PFA in PBS at room temperature and washed once with PBS. Cells were imaged by epi-fluorescence microscopy with excitation wavelength of 485 nm and emission wavelength of 600 nm using an Axioplan2 imaging system (Zeiss) coupled to a Cool SNAP HQ camera (Roper Scientific) with a Plan-NeoFluar 40×/1.3 objective. Image processing and fluorescence intensity measurements were performed using ImageJ software v1.53k ^43^.

### PTCH1 expression and purification

For PTCH1 purification, ExpiF293 cells were transiently transfected with polyethyleneimide (PEI) with the pCAGEN-FLAG-PTCH1-10His vector and grown in 200ml Expi medium 200ml in a 1L flask at 37°, 8% CO2 for 48 hours. Cells were pelleted by centrifugation and stored at -70°C.

pCAGEN-FLAG-PTCH1-10His cell pellets were thawed and resuspended in 16 mL of lysis buffer containing 25 mM Tris-HCl pH 8, 300 mM NaCl, 116 ng/mL benzonase, 1X PIC (Protease Inhibitor Cocktail 1000X: 1mg/ml aprotinin, leupeptin, pepstatin, antipain (Sigma Aldrich); 1M benzamidine in DMSO). Cells were lysed using a syringe and performing 10 back-and-forth strokes. The cell lysate was ultracentrifuged for 25 min at 100,000 g, at 4°C, using the TLA 100.3 rotor (Beckman). The pellet containing the membranes was recovered and homogenized in solubilization buffer containing 25 mM Tris-HCl pH 8, 300 mM NaCl, 1% dodecyl ß-maltoside (DDM, Anatrace), 0.2% cholesterol hemisuccinate, and 1X PIC. The mixture was incubated for 2 hours at 4°C with rotation, then ultracentrifuged for 25 min at 100,000 g at 4°C using the TLA 100.3 rotor. The supernatant containing the solubilized proteins was collected.

Anti-Flag resin (ANTI-FLAG M2 Affinity Gel, Sigma Aldrich) was washed with Mili-Q water and then equilibrated with purification buffer (25mM Tris-HCl pH8, 300mM NaCl, 0.0075% Lauryl Maltose Neopentyl Glycol (LMNG, Anatrace). The solubilized proteins were mixed with the anti-Flag resin and incubated for 90 min at 4°C with rotation. The sample was then introduced into a purification column at 4°C, and the resin was washed four times with purification buffer. The protein was eluted with 4 x 1 mL of purification buffer supplemented with 0.2 mg/mL flag peptide (Sigma Aldrich). The purification fractions were collected and loaded onto a 12% acrylamide SDS-PAGE gel.

The elution fractions showing a band of the PTCH1 protein at approximately 140 kDa (E1, E2, and E3) were pooled and used as such for affinity measurements. For structural studies, they were concentrated to 300 µL (100,000 MWCO PES, Sartorius Stedium) at 800 g and 4°C. The 300 µL of protein was injected into a Superose 6 increase 10/300 GL (Cytiva) size exclusion chromatography column on an Åkta Pure (Cytiva) device.

The fractions corresponding to the PTCH1 protein peak were pooled and then concentrated in small Amicon 100 kDa concentrators to 30 µL. The protein concentration was evaluated using a Nanophotometer UV-visible photometer (IMPLEN) and a 159850 mol^-1^. L. cm^-1^ extinction coefficient at 280nm.

### PTCH1-PAH Affinity measurements

Using microscale thermophoresis (MST): 30 µg/mL of membranes from HEK293-PTCH1 cells were incubated with 20 nM of tris-NTA-NT647 fluorescent probe (Nanotemper), and then with 250 µM to 15 nM of PAH. After a short incubation, the samples were loaded into capillaries (Nanotemper) in a MST analysis system (Monolith NT.115 (Nanotemper)) as described in ^37^. The dissociation constant (Kd) was calculated using the Nanotemper software.

#### Using intrinsic fluorescence variations

Measurements of the intrinsic fluorescence of PTCH1 tryptophan residues over time were performed in triplicate with an excitation wavelength of 295 nm and an emission wavelength of 330 nm using a spectrofluorimeter (Photon Technology International PTI - LPS-220). An initial measurement with 100 µl at 4.2 µM of PTCH1 purified after anti-Flag affinity purification was performed, then increasing concentrations of PAH were added, with 5 minutes of incubation between each measurement. The triplicate data were averaged, normalised and analysed using Kaleidagraph (Synergy). The formula y = FluoMax – [(FluoMax ™ FluoMin) * x / (Kd + x)] where x is the PAH concentration, Kd is the dissociation constant and y the measured fluorescence value was fitted using Kaleidagraph (Synergy Software).

### Cryo-em grid preparation, data collection and processing

PTCH1 at 1 mg/mL was incubated with PAH at a 1:50 PTCH1:PAH molar ratio for 30 minutes prior to grid preparation. Two applications of three micro-litres of purified PTCH1 incubated with PAH were applied to UltraAufoil (Quantifoil) 1.2/1.3 holey gold grids previously glow-discharged in air for 25s at 15 mA (Pelco Easiglow). A Vitrobot mark IV (Thermofisher) was operated at 10°C and 100% relative humidity with a waiting time of 7seconds, blot time of 3 seconds with blot force 0.

Grids were clipped and inserted in sample holder cassette for data collection in the cryo electron microscope. The SOLEIL synchrotron Polaris beamline (https://france-cryo-em.fr/) Titan Krios G4 (ThermoFisher) is a 300 kV column equipped with a cold FEG, a Selectris X energy filter, and a Falcon 4i electron detector. 26500 movies were collected at a magnification of 165000X and pixel size of 0.75Å with defocus ranging between -2 µm and -0.6 µm. The dose was 40 e-/Å^2^.

Data were processed using the Cryosparc V5 software ^44^. Movies were corrected using patch motion correction and processed with patch CTF option. Particle picking was done on a subset of micrographs using a CrYOLO particle picking ^45^ followed by 2D classification and selection to generate templates for Topaz training ^46^. Then 2-3 cycles of Topaz extract-2D classification-selection-topaz training were performed until convergence. Finally, Topaz extraction was performed on the full dataset. The particle set was sorted using 2D classification then several cycles of *ab initio* classification followed by non-uniform refinement.

### Model building and refinement

An Alphafold3^47^ model of human PTCH1 was fitted into the cryo-em map and adjusted with Coot version 0.98 software ^48^ and the Isolde^49^ plug-in to the ChimeraX^50^ interface. The PAH inhibitor geometry parameters were calculated with Phenix.elbow starting from atomic coordinates provided by Meliné Simsir (personal communication). The PTCH1-PAH complex was refined in real space using Phenix.RealSpaceRefinement ^51^. Model figures were done using ChimeraX version 1.10 ^50^ and Pymol 3.0 (Schrodinger). The channel was identified using the MoleOnline web server ^52^.

## Acknowledgements

We acknowledge SOLEIL synchrotron and the French EQUIPEX+ France Cryo-EM (ANR-21-ESRE-0046) for provision of cryo-EM facilities and we would like to thank Eric Larqué, Pierre Legrand, Heddy Soufari and Andrew Thompson for assistance in using POLARIS microscope.

We thank the Pasteur Institute Nanoimaging facility, the Curie Institute cryo-EM facility, the European Synchrotron Radiation Facility for provision of beam time on CM01 Titan microscope. We are grateful to E. Kandhia (ESRF), Daniel Levy, Aurélie Di Cicco and Julien Maufront (I. Curie) for assistance to electron microscopes. This work benefited from access to the EMBL Heidelberg imaging facility, an Instruct-ERIC centre and from assistance by Simon Fromm. Access to the Marseille eukaryotic cell culture facility was partly supported by the French Infrastructure for Integrated Structural Biology (FRISBI) ANR-10-INBS-0005. We received support from the ANR-22-CE11-0039 Oligo-ForceSensor project, ITMO Cancer equipment (**21CQ014-00)**, Département des Alpes Maritime (2020-461DGA-DSH), and CNRS Innovation. We acknowledge access to the Crystallography and Biophysics platform of the IBPC that is supported by the CNRS, the Labex DYNAMO (ANR-11-LABX-0011), the Equipex CACSICE (ANR-11-EQPX-0008) and the Conseil Régional d’Île-de-France (SESAME grant).

## Author contribution

OH performed protein purification and biochemistry experiments and collected cryo-EM data, MW produced HEK293T stably expressing FLAG-PTCH1-10His and performed the functional characterization of FLAG-PTCH1-10His over-expressed in these cells. CD produced Expi293 cells to express PTCH1. SA and SK prepared PAH. IMV designed and supervised the functional experiments of FLAG-PTCH1-10His in HEK293T cells and wrote the manuscript. VB designed and supervised the biochemistry and structural experiments, processed data and wrote the manuscript.

## Figure legends

Suppl. 1 cryo-em data collection and processing

---

**Supplementary figure S1.**
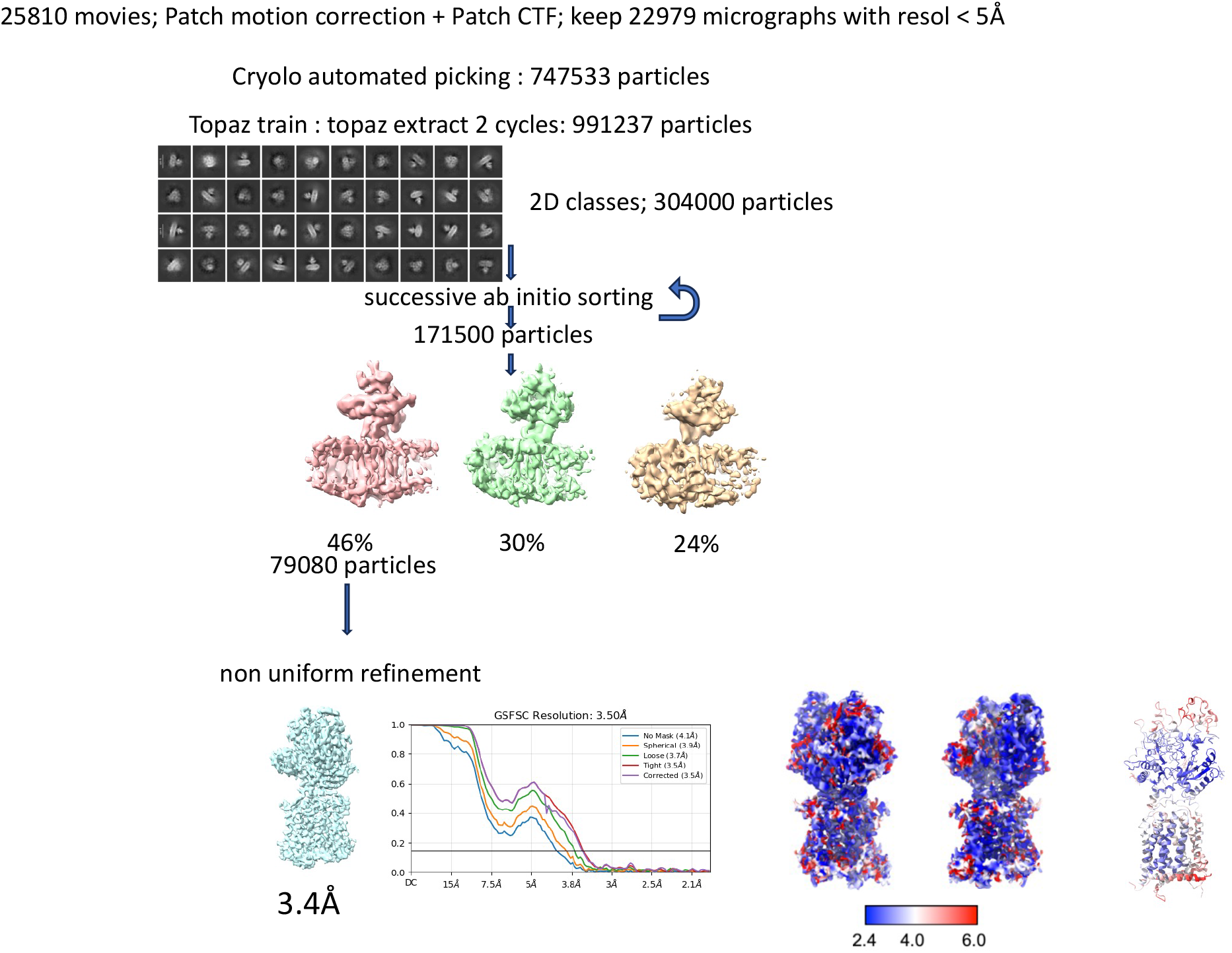
data processing with Cryosparc and Cryolo

